# SDRAP for annotating scrambled or rearranged genomes

**DOI:** 10.1101/2022.10.24.513505

**Authors:** Jasper Braun, Rafik Neme, Yi Feng, Laura F. Landweber, Nataša Jonoska

**Affiliations:** Department of Mathematics and Statistics, University of South Florida, Tampa, FL 33620, USA; Department of Chemistry and Biology, Universidad del Norte, Barranquilla, Colombia; Departments of Biochemistry and Molecular Biophysics, and Biological Sciences, Columbia University, New York, NY 10032, USA; Division of Clinical Pathology, Department of Pathology, Beth Israel Deaconess Medical Center, Boston, MA 02215

## Abstract

DNA rearrangements are important in various contexts, such as in vertebrate immunity, and cancer genome instability. The single-celled eukaryote *Oxytricha trifallax* undergoes massive and reproducible genome rearrangement during post-zygotic development, making it a compelling model organism to study DNA rearrangements. To date, computational tools for the extraction and analysis of rearrangement annotations lack transparency and rely on assumptions that may not hold for all analyzed data, leading to irreproducibility of results and loss of information through data filtering or misrepresentation.

An implementation of a procedure for the annotation and analysis of DNA rearrangement as a web application is discussed and tested. The resulting annotations provide an improvement over previous annotations in the following manner. (a) SDRAP achieves more complete precursor-product mappings than previous software (b) the software allows for full transparency of all parameters used during the annotation and therefore facilitates reproducible results, and (c) this parameter transparency makes SDRAP suitable for comparison of genomic data from different sources, including cancer genomes.

This work introduces a theoretical framework and software to systematically extract and analyze annotations from pairs of genome assemblies corresponding to precursor and product rearrangement data. The software makes no assumptions about the structure of the rearrangements, and permits the user to select parameters to suit the data.

## 1 Background

Genome sequences sometimes undergo large-scale rearrangements that alter the order and content of the DNA [11, 18, 22]. These events can impact organisms in highly diverse ways and on diverse timescales, ranging from chromosome inversions or translocations that contribute to species and population diversity [10, 22], to the evolution of adaptive immune systems [19], tissue differentiation [20], or nefarious cancers [26]. In some eukaryotes, such as the ciliate *Oxytricha trifallax*, programmed genome rearrangements permit massive structural modifications from a precursor genome (the germline micronucleus) into a product genome (the somatic macronucleus) [8, 25]. As a result, these two genomes, though housed in the same cell, are left with reduced similarity to each other, permitting comparative genomics within a single cell. The rearrangement process includes removal of more than 90% of the precursor DNA [8], and reorganization of the remaining DNA fragments into ∼ 18,000 different gene-sized nanochromosomes as the product [14]. The order or orientation of the DNA segments in the precursor is often different than in the product [17]. The presence of such complex DNA rearrangements during macronuclear development makes *O. trifallax* and other species of ciliates powerful model organisms to study DNA rearrangement processes that appear in a wider range of organisms. For example, V(D)J recombination in vertebrates also entails genome rearrangements [19], and somatic rearrangements are commonly associated with certain types of genome instability in cancers, including the widespread phenomenon of chromothripsis [9].

Existing models of DNA rearrangements apply only to one-to-one, or one-to-many mappings [3, 4, 13, 12, 23], but in the context of ciliate genomics, a many-to-many correspondence between regions in rearrangement precursor and product is observed. Previous approaches to annotating ciliate genomes have been performed in species-specific ways, using custom scripts, with many complex cases left untreated, and many annotation parameters chosen arbitrarily without a precise definition of the parameters associated with the output [7, 8, 24]; these approaches complicate the reproducibilty of annotations, prevent crossspecies comparisons, and lack transparency and comprehensiveness. Here, we present a protocol that readily describes and outputs relationships between a pair of precursor and product genomes. The algorithm also annotates positions of telomere addition sites, specifies the types of arrangements between precursor and product loci, and detects inversions or translocations among those arrangements (scrambled maps). We present an implementation of our algorithm called *Scrambled DNA Rearrangement Annotation Protocol (SDRAP)* that allows a range of parameters, such that many genomes with similar general properties to each other but specific differences can be accommodated and processed with this suite. Further, we specify definitions related to retained and eliminated DNA sequences, scrambling, paralogy and completeness of loci, which offer improvements on existing models of genome rearrangements. Together, this annotation tool allows consistent, automated, adaptable and reproducible analysis of scrambled pairs of genomes, such as the rearrangements that occur during somatic differentiation in ciliates or other cell types.

We begin by defining the notions used in SDRAP (2). These include a match as a pair of intervals that have ‘matching’ DNA between the precursor and the product, arrangements as sets of matches corresponding to the same product contig, and types of unambiguous arrangements. In 2.4 we describe alignment adjustments that have to be performed after BLAST that are pertinent to situations of a scrambled genome. The algorithm describing the steps of the protocol is described in 2.5 and the implementation is discussed in 3. We conclude with results of the implementation and a short discussion.

## 2 Preliminaries

### 2.1 Annotation of DNA Rearrangements

DNA rearrangements considered here involve reordering and/or inversion of segments from a precursor sequence to form a product sequence. In the context of ciliate biology for example, precursor and product sequences represent portions of the micronuclear and macronuclear chromosomes, respectively. Limitations of sequencing technologies, as well as read assembly and scaffolding algorithms, result in a fragmented assembly of the precursor genome for the ciliate *O. trifallax* [8] which is a model organism to study various genomic processes. Product chromosomes, on the other hand, are relatively short (mostly one to two genes, not more than 8 genes on a chromosome) and in recent efforts to improve the macronuclear genome assembly of the ciliate *O. trifallax*, these chromosomes were sequenced using long reads that frequently cover entire chromosomes [14].

### 2.2 Matches and Arrangements

The rearranging segments in precursor and product represent sequence homology (i.e., local sequence alignments), and their rearrangement is fully described by the locations of the segments in the respective genome assemblies. Hence, the start and the end coordinates of the segments (without specifying the underlying DNA sequence) can represent the rearrangement. Hence a pair of corresponding precursor and product segments is represented by a triple *M* = ([*a, b*], [*c, d*], *σ*), called a **match**, where [*a, b*] is an integer interval called **precursor interval** (denoted Prec(*M*)) that describes the location of the segment in the precursor, and [*c, d*] is an integer interval called **product interval** (denoted Prod(*M*)) that describes the location of the segment in the product, while *σ* ∈ {0, 1} is the **orientation** of the match, denoted *σ*(*M*) (note that the term “match” here does not refer to a pair of matching residues in a sequence alignment as commonly expressed in the literature). We follow the convention that *σ* = 1 indicates that the two segments are found in the same orientation, and *σ* = 0 indicates that the two segments are found in opposite orientations of the respective precursor and product sequences. Local sequence alignments can be viewed as matches in the obvious way, and the language introduced here for matches is often also used for alignments.

To determine whether or not segments have to be permuted to form a product sequence, we must be able to compare order and orientation of precursor intervals with corresponding product intervals. A natural ordering on a set of intervals can be defined by:

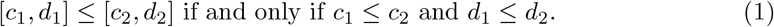

However, this ordering is a total ordering only on sets in which no interval is contained in another. Thus, we are interested in sets of matches that satisfy:

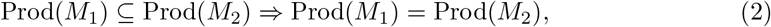

for all matches *M*_1_, and *M*_2_ in a given set. In this way distinct product intervals of the matches in the set are never contained in one another. A set of matches that satisfies (2) is called an **arrangement**. Hence, product intervals in an arrangement ℳ are totally ordered. We observe that a subset of an arrangement is also an arrangement.

To each match *M* in an arrangement ℳ, we assign an index *i*_ℳ_ (*M*) that indicates the position of Prod(*M*) in the order of ℳ. We call *i*_ℳ_ (*M*) the **index of** *M* **in** ℳ. Observe that the condition (2) references the product interval only, hence in the following we describe unambiguous arrangements that arise according to the order of precursor intervals.

### 2.3 Properties of Arrangements

The ordering of the product intervals of an arrangement defined by (1) can contain an ambiguous correspondence between precursor and product intervals established by the matches.

This situation prevents a comparison of the ordering of the precursor and product segments. For example, the same precursor region may correspond to two regions in the product, so that a one-to-one correspondence between precursor and product segments may not exist (such as matches *M*_4_ and *M*_5_ in Figure 1(a)). Such overlap of precursor intervals in an arrangement prevents the interval ordering defined with (1) to be an (total) ordering on the set of precursor intervals of an arrangement, in particular when one precursor interval contains another. The general strategy taken in the algorithm below is to consider all maximal subsets of an arrangement that do not have these issues.

**Figure 1:**
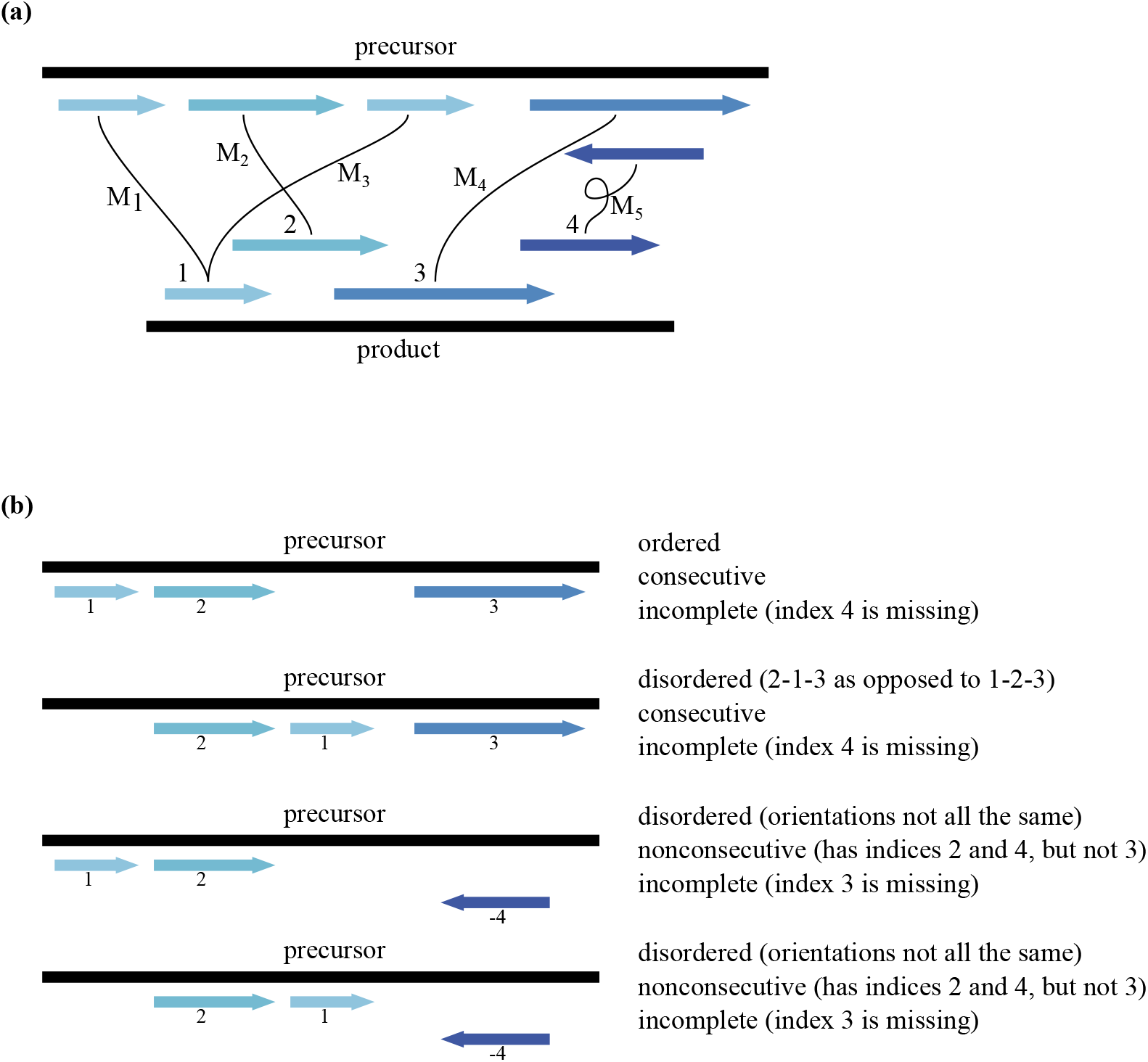
**(a)** An arrangement of matches {*M*_1_, …, *M*_5_} with their indices labeling the product intervals. Matches *M*_1_ and *M*_3_ share a product interval and are given the same index 1. A precursor and a product interval connected by a black line form a match. An inverted match is indicated by the loop in the black line and reverse oriented arrows. The black lines are labelled by the indices of the corresponding matches. **(b)** Unambiguous subarrangements of the arrangment in (a). Only the precursor intervals are shown from the top figure. The index is negated for the reverse oriented match. For each of the unambiguous subarrangements, it is indicated whether or not the arrangement is ordered, consecutive, or complete. There is no complete unambiguous subarrangement because Prec(*M*_5_) ⊆ Prec(*M*_4_).

Given a positive integer *p*, two matches *M*_1_, *M*_2_ are considered *p***-overlapping** if:

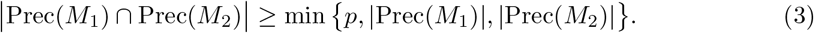

A subset ℳ′ of an arrangement ℳ is **unambiguous** if:

i. *i*_ℳ_ (*M*_1_) ≠ *i*_ℳ_ (*M*_2_), for all *M*_1_, *M*_2_ ∈ ℳ′, and
ii. No two matches in ℳ′ are *p*-overlapping, and
iii. ℳ′ is a maximal set of matches with the first two properties.

We consider an unambiguous set of matches **ordered** if either they all have orientation *σ* = 1 and appear in the same order on the precursor as on the product, or if they all have orientation *σ* = 0 and they appear on the precursor in reverse order compared to the product (see Figure 1(b)).

An unambiguous subset ℳ′ of an arrangement ℳ is called **consecutive** if the set of indices ℐ _ℳ_ (ℳ′) = {*i*_ℳ_ (*M*) : *M* ∈ ℳ′} forms a consecutive set of integers, and it is called **complete** with respect to ℳ if the set of indices ℐ _ℳ_ (ℳ′) is the same as the set of indices ℐ _ℳ_ (ℳ′) = {*i*_ℳ_ (*M*) : *M* ∈ ℳ} of matches in. An unambiguous arrangement ℳ is not scrambled if it is ordered, consecutive and complete. Given a set of properties 𝒮 ⊆ {ordered, consecutive, complete}, an unambiguous subset of an arrangement is called 𝒮-scrambled if one or more of the properties in 𝒮 is violated.

To assess whether an arrangement ℳ is scrambled, we consider all possible unambiguous subsets of ℳ. Whenever at least one of the unambiguous subsets of ℳ is 𝒮-scrambled, the parent set ℳ is called **weakly 𝒮-scrambled**. If all non-singleton unambiguous subsets of are 𝒮-scrambled, the parent set is called **strongly 𝒮-scrambled**. Weak and strong versions of the properties ordered, consecutive, and complete are defined analogously. Here, the terms weak and strong refer to the strength of the available evidence that designates an arrangement as scrambled, such that the user can decide the level of stringency whenever the feature “scrambled” becomes biological or statistically relevant.. Figure 1 shows an example of an arrangement together with its unambiguous subarrangements. With 𝒮 = ordered, consecutive, the depicted arrangement is weakly, but not strongly 𝒮-scrambled. However this set is strongly scrambled because there is no complete unambiguous subarrangement.

### 2.4 Merging Alignments

Since efficient and well-established sequence alignment tools (like BLAST [1, 2]) for the identification of the rearranging precursor and product segments already exist, our protocol starts with a set of local sequence alignments. In the context of DNA rearrangements in ciliates, high sequence similarity between corresponding precursor and product segments can be expected. Some insertions, deletions and substitutions can be attributed to allelic variation, sequencing errors, and ambiguities during read assembly. Thus, gapped local sequence alignment algorithms with high sequence similarity that identify precursor and product segments participating in the rearrangement are prefered.

Standard gapped sequence alignment algorithms, including the ‘gapped’ option in BLAST, may mistake strings of non-scrambled product segments with short overlap being within short distances in the precursor as single long gapped alignments. However, such situations appear in scrambled genomes and should be represented as multiple consecutive matches. Thus, a high degree of control over the introduction of gaps in the sequence alignments is necessary. One strategy taken in the past has been to instruct BLAST to search for ungapped alignments and then to further process these alignments [7, 6, 8]. We adopt this general strategy in our software and provide complete user control for choosing parameters that combine ungapped alignments into longer gapped alignments.

Informally, the algorithm merges two ungapped alignments when they have the same orientation, their precursor and product intervals are not ‘too far’ apart, and the gap that must be introduced to connect the two alignments is not ‘too large’. We specify these notions below.

Given two equally oriented matches *M*_1_ = ([*a*_1_, *b*_1_], [*c*_1_, *d*_1_], *σ*) and *M*_2_ = ([*a*_2_, *b*_2_], [*c*_2_, *d*_2_], *σ*), the **merge** of *M*_1_ and *M*_2_ is:

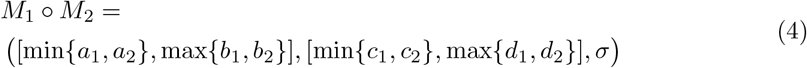

In the case when Prod(*M*_1_) ≤Prod(*M*_2_) is the match and Prec(*M*_1_) ≤ Prec(*M*_2_) the merge of *M*_1_ and *M*_2_ *M*_1_ ∘ *M*_2_ = ([*a*_1_, *b*_2_], [*c*_1_, *d*_2_], *σ*).

We constrain the conditions by which the two matches can be merged to form a larger alignment that spans the combined precursor and product regions with two constraints, shift and distance. The **start shift** of two matches to be

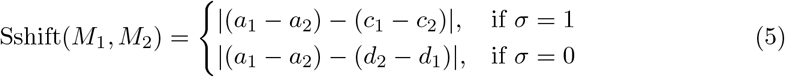

and the **end shift** for two matches is

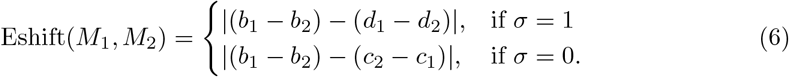

In Figure 2(a) the start shift is indicated with purple and the end shift is shown in red. The overall **shift** of two matches is then:

**Figure 2:**
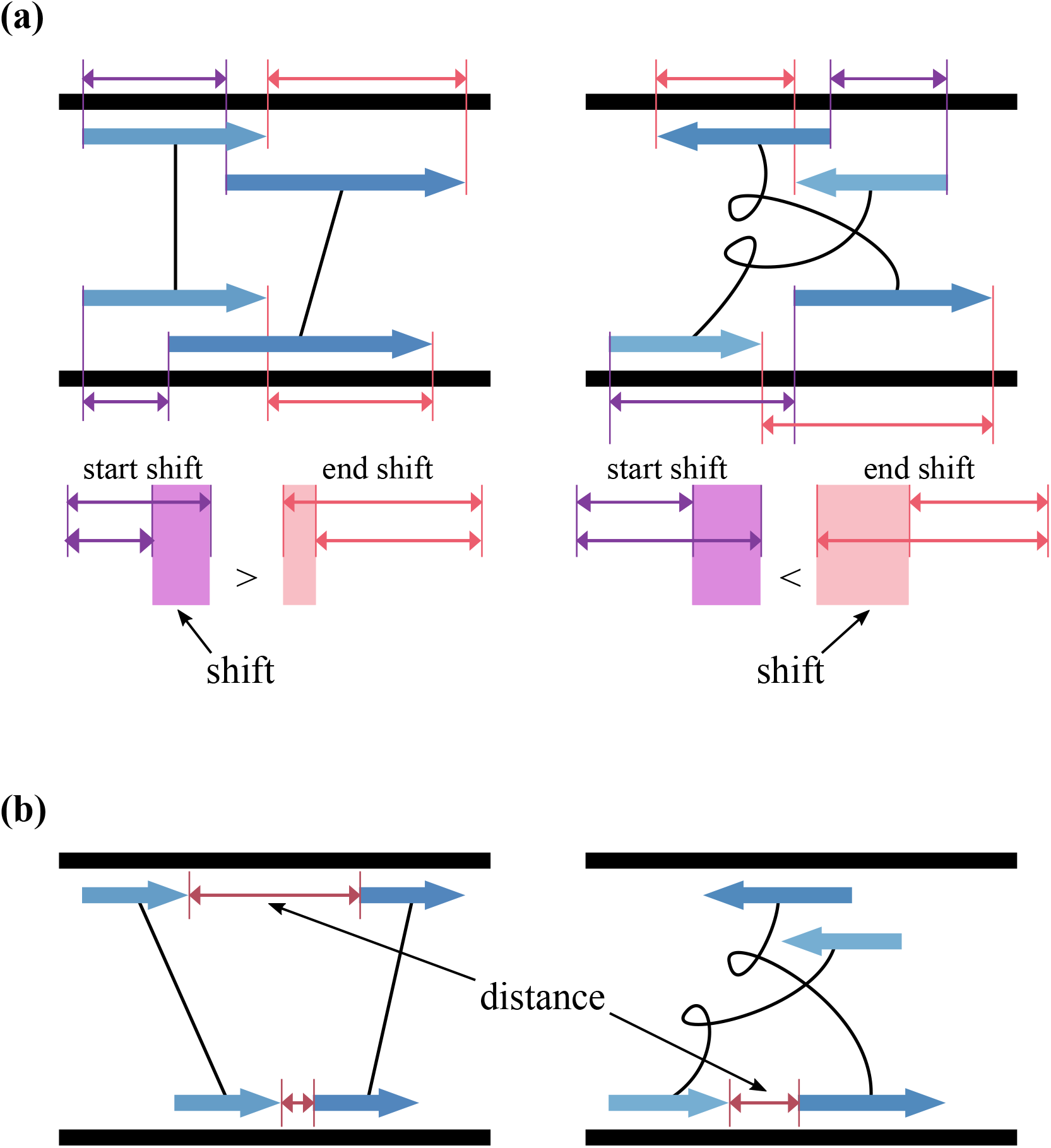
**(a)** Illustration of the shift between two matches with orientation *σ* = 1 (left) and orientation *σ* = 0 (right). The start shift is depicted in purple and the end shift is depicted in red. The larger of the start and end shifts is the shift between the two matches. **(b)** Illustration of the distance between two matches.

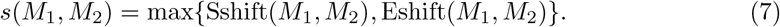

Figure 2(a) depicts examples of the shift between two matches. When *M*_1_ and *M*_2_ are ungapped alignments, their shift is precisely the size of the gap that must be introduced to form the alignment *M*_1_ *M*_2_. Note that the shift between two matches conveys no information about the gap or overlap between their precursor and product intervals.

Define the **distance** between *M*_1_ and *M*_2_ as:

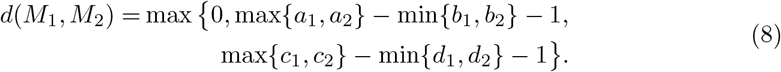

The distance between *M*_1_ and *M*_2_ is the larger of the spaces between their precursor intervals and between their product intervals. In the case when Prod(*M*_1_) ≤ Pr{od(*M*_2_) and Prec(*M*_1_) ≤ Prec(*M*_2_) the distance between *M*_1_ and *M*_2_ is *d*(*M*_1_, *M*_2_) = max {0, *a*_2_ − *b*_1_ − 1, *c*_2_ − *d*_1_ − 1} (see Figure 2(b)).

Finally, two matches *M*_1_ and *M*_2_ with the same orientation are (*s*_max_, *d*_max_)-**mergeable** if:

i. *σ*(*M*_1_) = *σ*(*M*_2_), and
ii. *s*(*M*_1_, *M*_2_) ≤ *s*_max_, and
iii. *d*(*M*_1_, *M*_2_) ≤ *d*_max_.

Observe that *s*_max_ directly controls the maximum gap or overlap that may be introduced between two ungapped matches to be merged. While *d*_max_ bounds how far apart aligned regions of the two matches that are merged may be on the precursor or the product sequence.

### 2.5 Annotation Protocol

The algorithm takes as input a set ℋ of ungapped local sequence alignments between a precursor and product sequence (in our implementation these are produced by BLAST with ungapped option) and returns arrangements ℳ with information whether each ℳ is weakly and strongly 𝒮-scrambled. An arrangement ℳ consists of matches that represent members of ℋ directly, or are obtained by merged members of ℋ. The input set ℋ may not satisfy requirement (2), so some members of ℋ would be excluded in production of ℳ. Informally, the procedure can be broken down into 3 steps as explained in 2.5.1, 2.5.2, and 2.5.3, respectively:

1. Extract ‘preliminary matches’ from H for which condition (2) is enforced (the set ℳ_pre_).
2. Add additional, potentially paralougus matches from the remaining alignments that may violate condition (2).
3. Determine whether or not the set of matches obtained in the first two steps is weakly and/or strongly 𝒮-scrambled.

We describe in details these three steps.

#### 2.5.1 Preliminary Matches

In order to ensure property 2 holds and to reduce the redundancies in the matches, we enforce non-containment of product intervals in a preliminary arrangement annotation step. In this step, we exclude alignments whenever their product intervals are already covered by product intervals of matches already in the current set.

Given a positive integer *c*, a match *M* is *c***-covered** by a set of matches ℳ, if:

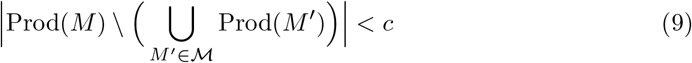

Informally, *M* is *c*-covered by ℳ if Prod(*M*) has fewer than *c* base pairs that are not ‘covered’ by members of ℳ. In other words, Prod(*M*) is considered covered by the product intervals of other matches in ℳ if less than *c* base pairs of *M* are not covered by their union. We denote the expression on the left-hand side of (9) by ncov(*M*, ℳ), or ncov(*M*, ℳ′), when ℋ = { *M*′}. Given the set of potential matches ℋ produced by the ungapped option of BLAST, a subset ℳ_pre_ of ℋ is obtained such that no match *M* ∈ ℳ_pre_ is *c*-covered by its complement ℳ_pre_ \ {*M* }. The set ℳ_pre_ is the set of **preliminary matches**. This set is obtained by sorting the input H by alignment bitscore. An alignment *H* of ℋ is added to ℳ_pre_ unless *H* is *c*-covered by the current set ℳ_pre_. In this process it is tested whether *H* is (*s*_max_, *d*_max_)-mergeable with any of the current members of ℳ_pre_, and when merge is performed, those members of _pre_ that are *c*-covered by *H* are removed. A flowchart and an example for this process are given in Figure 3.

**Figure 3:**
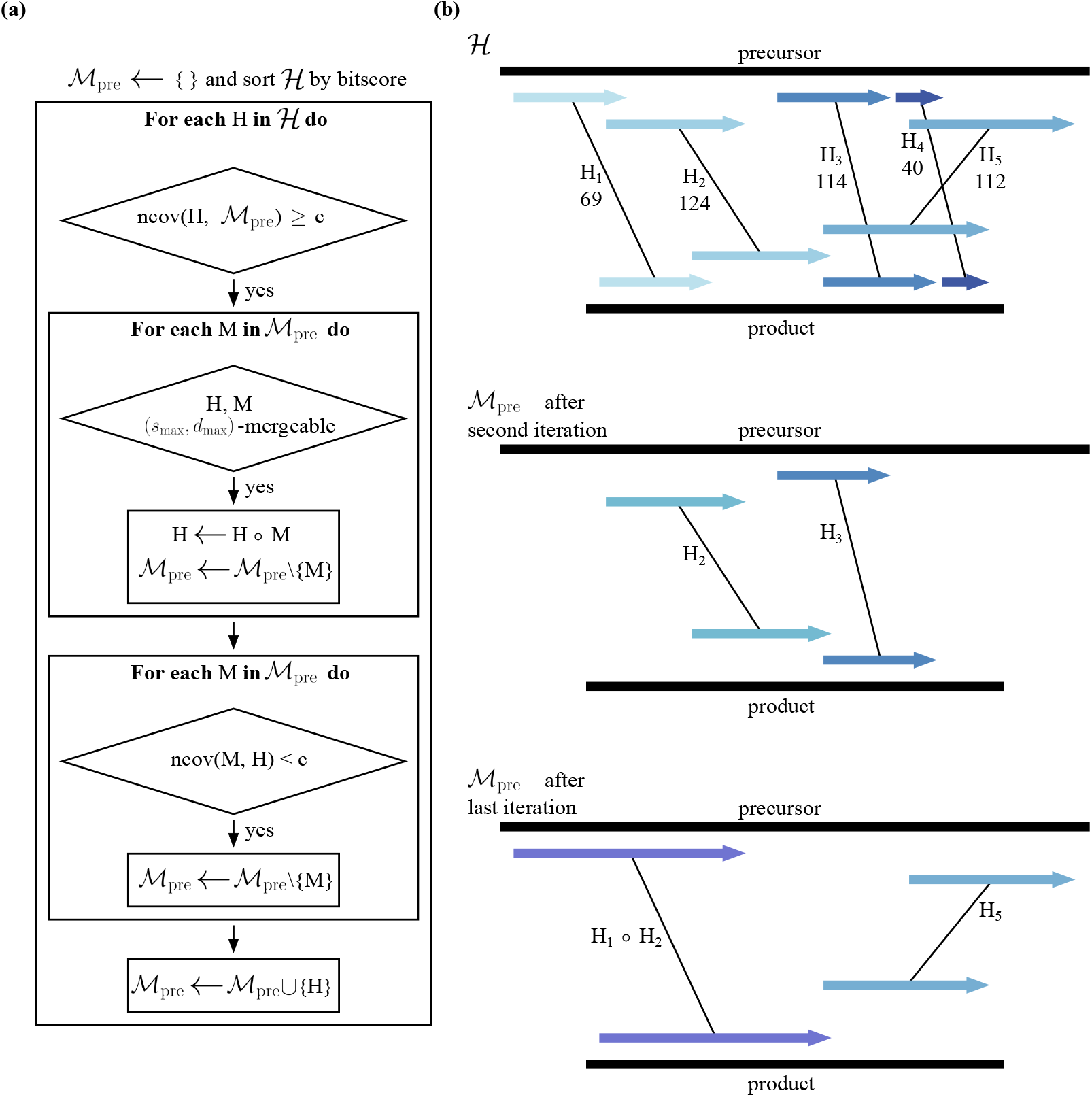
**(a)** A flowchart for the preliminary arrangement annotation algorithm. In **(b)**, the input set ℋ = {*H*_1_, …, *H*_5_} is shown at the top with numbers below their labels indicating their bitscores. The bitscore orders ℋ to (*H*_2_, *H*_3_, *H*_5_, *H*_4_, *H*_1_). The set ℳ _pre_ after the second and last iterations of the outer loop in the flowchart from (a) is depicted in the middle and bottom, respectively. The first two iterations form matches from *H*_2_ and *H*_3_ and add them to ℳ _pre_ without merging. *H*_3_ is removed after addition of *H*_5_ because *H*_5_ covers *H*_3_. For the same reason *H*_4_ is not added to ℳ _pre_. The last addition of *H*_1_ merges *H*_1_ and *H*_2_. At the end of this algorithm, ℳ _pre_ = {*M*_1_ = *H*_1_ ∘ *H*_2_, *M*_2_ = *H*_5_} forms an arrangement and indices can be assigned to its members 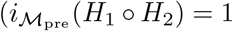 and 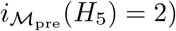.

Observe that since *c >* 0, testing for *c*-coverage procedure guarantees satisfaction of (2). The parameter *c* allows for leniency in the precision by which (2) is enforced for preliminary matches. A product segment may match multiple regions in the precursor, but with some discrepancies at the region’s boundaries. The parameter *c* helps controlling how strict the algorithm should view product region boundaries.

The final step of this part of the protocol is assignment of indices to each match in ℳ_pre_.

#### 2.5.2 Additional Matches

Given a real number *r* ∈ [0, 1], an alignment *H*, and a preliminary match *M* in ℳ_pre_, we say that *H* **intersects** *M* **sufficiently** if:

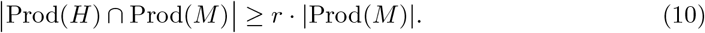

In the second step of the procedure, a set ℳ_add_ of additional matches are derived from the set ℋ′ ⊆ ℋ \ ℳ_pre_. We exclude alignments from H that contributed to set ℳ_pre_. Similarly as in the first step constructing ℳ_pre_, for each alignment *H* ∈ ℋ′, the algorithm merges *H* with members of the current set of additional matches ℳ_add_, whenever appropriate, and then compares it to each of the preliminary matches. Whenever the intersection of the (possibly merged) match *H* with a member *M* ∈ ℳ_pre_ is sufficiently large, *H* is added to ℳ_add_ and viewed as having the same index as *M* in _pre_. A flowchart and an example for this part of the algorithm are given in Figure 4.

**Figure 4:**
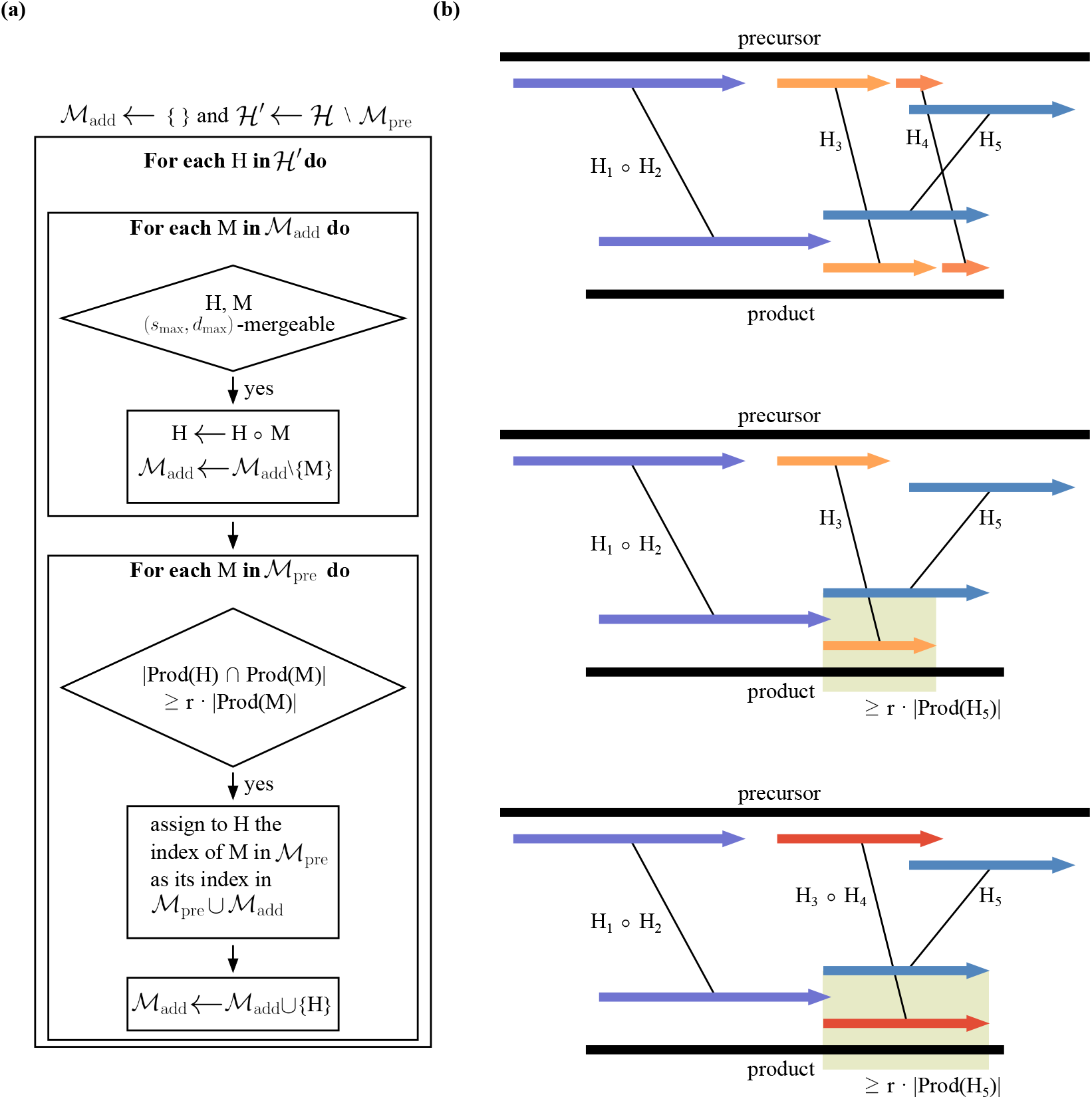
**(a)** A flowchart for the additional match annotation algorithm. The assignment ℋ′ ← ℋ \ ℳ _pre_ indicates that members from ℋ that form members of ℳ_pre_ either directly, or indirectly via merging, are not included in ℋ′. The example in **(b)** continues the example from Figure 3(b). Alignments *H*_3_ and *H*_4_ are in H′. In the first iteration *H*_3_ is added to ℳ _add_. In the second iteration, because *H*_3_ and *H*_4_ are (*s*_max_, *d*_max_)-mergeable, *H*_3_ ∘ *H*_4_ replaces *H*_3_ in ℳ _add_. At the end of this algorithm, because *H*_5_ and *H*_3_ ∘ *H*_4_ cover the same product interval, the match *H*_3_ ∘ *H*_4_ has index 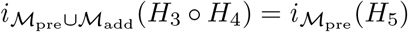. *H*_3_ ∘ *H*_4_ and *H*_5_ may represent paralogues in the precursor.

The parameter *r* determines how much the product interval of an alignment must overlap with the product interval of a preliminary match i order to inherit its index. Such an additional match inheriting the index of a preliminary match *M* can be viewed as a repeated region in the precursor that matches the same region in the product and detects possible paralogues in the genome. Conceptually, for detection of repeats in the genome, the value of *r* should be close to 1 for the two matches. Note that a single alignment *H* may give rise to multiple additional matches (with distinct indices) during the additional match extraction step if its intersection with multiple preliminary matches is sufficiently large. However, since preference is given to alignments with higher bitscore in the first step of the procedure, and longer alignments are statistically more significant, duplicate appearances of alignments in ℳ_add_ are rare. When the input alignments have high percent identity, such double-counted alignments should be even rarer. On the other hand, when the percent identity of alignments in ℋ is not always high, then a long alignment may have less significance then each of the non-mergeable groups of alignments that collectively cover the product interval of the longer alignment. In this situation, the short alignments would make it into the set ℳ_pre_ whereas the longer alignment would be excluded from ℳ_pre_. Instead, the long alignment would give rise to multiple members in ℳ_add_, one for each of the smaller alignments that formed preliminary matches.

Below we somewhat abuse the notion of arrangement and we call the set ℳ = ℳ_pre_ ∪ ℳ_add_ an arrangement although the condition (2) may be violated with addition of M_add_. We are interested in the maximal unambiguous subarrangements of ℳ.

#### 2.5.3 Scrambling

In order to determine whether the arrangement ℳ= ℳ_pre_ ∪ ℳ_add_ is weakly or strongly 𝒮-scrambled, the algorithm must test whether the unambiguous subsets of ℳare 𝒮-scrambled. Therefore the maximal unambiguous subsets need to be extracted. To identify these sets, we associate a graph with a set of matches ℳ, by letting the matches be the vertices and two matches are connected with an edge whenever the two matches do not share the same index and are not *p*-overlapping. Then the maximal cliques in the graph correspond precisely to the maximal unambiguous subarrangements. The graph associated with the arrangement depicted in Figure 1 as well as the cliques corresponding to its unambiguous subarrangements is shown in Figure 5.

**Figure 5:**
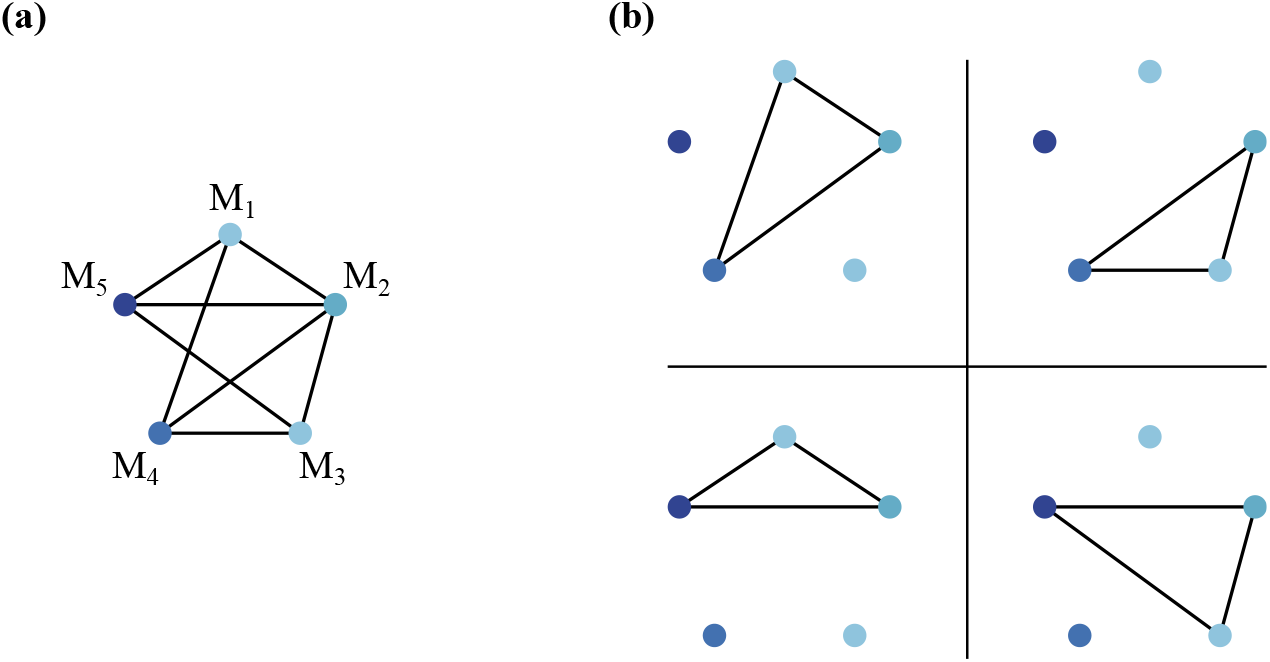
**(a)** The graph associated with the arrangement shown in Figure 1(a). **(b)** The maximal cliques of the graphs. The maximal cliques correspond to the unambiguous subarrangements in Figure 1(b).

While a set of matches theoretically can have an exponential number of unambiguous subsets with respect to the number of matches, in practice this number is much lower in most cases. For that reason, an output-sensitive algorithm to list maximal cliques, such as the algorithm introduced by [15], is included in SDRAP. Once an unambiguous subarrangement is detected, the procedure determines whether that subarrangement if weakly or strongly 𝒮-scrambled by checking the properties of the corresponding index sets.

## 3 Implementation

The algorithm described in 2.5 was implemented as a web application called **Scrambled DNA Rearrangement Annotation Protocol**, or **SDRAP**, using PHP 5.3.3 and MySQL 5.6.31 with Apache 2.2.15 on a linux server running with the CentOS 6.7 operating system. The user interface was implemented using HTML, CSS and javascript and can be accessed at https://knot.math.usf.edu/SDRAP. The code and a complete documentation for SDRAP are available at https://github.com/JasperBraun/SDRAP. The application accepts the input genome assemblies in FASTA format. The steps of the procedure are:

1. Detect and mask telomeric sequences at the ends of product sequences (identifying chromosome ends).
2. BLAST product sequences as query against precursor sequences as subject to obtain the set of alignments ℋ.
3. Annotate arrangements (form set ℳ = ℳ_pre_ ∪ ℳ_add_ from the set ℋ).
4. Compute arrangements (determine unambiguous subarrangements and their properties).

SDRAP implements its own telomere detection algorithm which is a heuristic adaptation of the Smith-Waterman gapped local sequence alignment algorithm [21]. Since many such variants exist which use the same or a similar strategy (such as BLAST and LFASTA), the algorithm is not discussed here, but a precise description can be found at https://github.com/JasperBraun/SDRAP. BLAST is run internally after telomeres are masked as the second step of the procedure. SDRAP uses Nucleotide-Nucleotide BLAST 2.2.31+ and the parameters are listed in the github link. For the remainder of the computation, SDRAP only considers the portions of the product sequences between the detected telomeres at either end, if any. When a telomere at the 5’ or 3’ end is missing, the product sequence included in subsequent steps includes the entire prefix or suffix, respectively.

SDRAP applies the arrangement annotation algorithm discussed in 2.5.1 and 2.5.2 to the set of local alignments returned by BLAST during the second step. The procedure returns annotations of precursor and product intervals of the resulting matches. In addition to the parameters *s*_max_, *d*_max_, *c* and *r* discussed in 2.5.1 and 2.5.2, the software accepts minimum thresholds 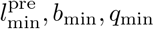 for length, bitscore and percent identity that alignments returned by BLAST must satisfy to be included in the set ℋ of alignments used for preliminary match annotation. Another set of parameters 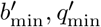 is provided which define minimum bitscore and percent identity thresholds that alignments must satisfy to be included in the set ∪′ of alignments used for additional matches.

Precursor and product interval annotations are complemented with annotations of pointers, gaps, **fragments** and eliminated sequences, which are defined here. Let ℳ_pre_ be the preliminary arrangement of a precursor and a product sequence. A **pointer** is the region of overlap, if any, between two product intervals with consecutive indices of preliminary matches, together with the corresponding two regions in the precursor. Since no two alignments in the preliminary arrangement ℳ_pre_ determined by the procedure in 2.5.1 have the same product interval, the overlapping region between two successive product intervals of preliminary matches *M*_*i*_, *M*_*j*_ corresponds to exactly two regions in the precursor sequence. These two regions are located at one of the ends of Prec(*M*_*i*_) and Prec(*M*_*j*_). SDRAP returns pointer annotations for both precursor and product. The **gaps** in a product sequence with respect to a precursor sequence are the regions in the product complementary to the product intervals of the preliminary matches. Gaps represent regions that are not covered by the precursor sequence, i.e., gaps in coverage. The regions on the precursor sequence in between precursor intervals of the union of all arrangements are annotated as **eliminated sequences**. Eliminated sequences represent the regions in the precursor that do not contribute to the product DNA. Minimum length bounds 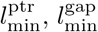, and 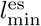 define the minimum lengths of pointers, gaps, and eliminated sequences, respectively, to be annotated as such. When one of these features is shorter in length than its respective bound, it is not annotated. All three bounds are provided as input parameters. Fragments of matches are those (possibly merged) matches in the set ℋ⌂ that are not returned as additional matches at the end of the additional match annotation algorithm described in 2.5.2. A match *H* in ℋ′ is not included in the additional match annotation if for all preliminary matches *M*, | ∩ Prod(*H*) ∩ Prod(*M*) | */* | Prod(*M*) |*< r*. All such matches *H* are annotated as fragments. Fragments represent statistically significant alignments that were not captured by the annotation because they are redundant to larger or higher scoring alignments. In order to annotate fragments with some specific overlapping properties, we check for inequality (10) for some threshold *r*′ *< r*. Whenever a fragment satisfies the inequality (10) for some preliminary match *M* and threshold *r*′, then a copy of it is annotated as a fragment with the index of *M*. When a fragment does not satisfy inequality (10) (with a lower threshold *r*′) for any preliminary match *M*, it is annotated as a fragment without index. The threshold *r*′ is provided as another input parameter of the software. All annotations are given labels providing as much information as possible to establish the correspondence of the annotated features in one genome assembly with location of corresponding annotations in the other.

To limit extraction of potentially exponential numbers of unambiguous subsets for each of the arrangements, SDRAP accepts an input parameter *k*_max_ which determines the maximum number of unambiguous subsets extracted from each arrangement. A table listing each arrangement and a number of properties of the arrangement is returned by SDRAP. The properties given for each arrangement include coverage of the product sequence by the matches in the arrangement, presence of gaps, presence of *p*-overlapping matches, presence of matches with the same index, and weak and strong 𝒮-scrambledness.

### 3.1 Algorithm Complexity

Since SDRAP is applied to potentially large datasets, the procedure’s computational complexity must be considered. The following complexity analysis can be broken up into two parts: (1) the arrangement annotation step and (2) the scrambling property computation step.

For the annotation of arrangements, alignments obtained using BLAST and derived matches can be represented as pairs of integers together with a binary orientation. Determining mergeability and subsequently merging two matches requires a constant number comparisons of orientation, and coordinates of precursor and product intervals. Calculating ncov(*M*, ℳ) for a match *M* and a set of matches ℳ requires iteration over ℳ with a constant check for how far the product interval of each member overlaps with that of *M*. Thus, the algorithm for extracting preliminary matches ℳ _pre_ runs through each *M* ∈ ℳ _pre_ for every *H* and therefore it takes *O*(*n*_pre_*h*) time, where *n*_pre_ = | ℳ _pre_ | and *h* = | ℋ |. This algorithm is applied to all of *N* precursor-product sequence pairs which have high-scoring pairs between them. After the set ℳ _pre_ is determined, assigning indexes 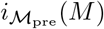 for each *M* ∈ ℳ_pre_ requires sorting and subsequent iteration over ℳ _pre_, totalling *O*(*n*_pre_ log *n*_pre_) operations. Next, to extract ℳ _add_ from the set ℋ′, the sets ℳ _pre_ and ℳ _add_ are iterated once for each *H* ∈ ℋ′ (see Figure 4), performing a constant number of iterations. Since ℋ′ ⊆ ℋ, and with *n*_add_ = | ℳ _add_ |, the additional arrangement annotation step has complexity *O*(*h*(*n*_pre_ + *n*_add_)).

To determine which arrangements are scrambled, a graph is associated with each arrangement (see Figure 5). The arrangement’s *n* = *n*_pre_ + *n*_add_ matches are represented by vertices and adjacency is determined via pairwise comparison. Unambiguous subarrangements correspond to maximal cliques in the graph and so determining all unambiguous subarrangements reduces to the problem of listing all maximal cliques. In general, a graph may have an exponential number (in the number of vertices) of maximal cliques. To avoid exponential complexity, a threshold *k*_max_ which limits the number of cliques to extract from each arrangement is imposed and the output-sensitive algorithm from [15], which scales as the cube of the number of extracted cliques, is applied. There are at most *n* connected components, which can be obtained using a simple *O*(*n* + *m*) DFS traversal, where *m* is the number of edges. Then extracting at most *k*_max_ cliques has complexity *O*(*n*^4^*k*_max_). The matches in the overall arrangement can be sorted according to product start coordinate prior to extracting cliques in *O*(*n* log *n*) time, so that subsequently determining arrangement properties of each of the unambiguous subarrangements corresponding to the cliques takes simple iteration over the matches. Since *k*_max_ is an input parameter, computing arrangement properties takes *O*(*n*^4^) operations.

## 4 Results and Discussion

Many parameter values of the software SDRAP should be chosen on a context-specific basis. To assess the effect different parameter values have on the computation, the procedure was run with various parameter values on the precursor and product genomes of the organism *Oxytricha trifallax*, obtained from [8] and [24], respectively. An outdated assembly of the macronuclear genome was chosen so that the results can be compared with previous annotations. In general, a low, a center and a high value was chosen for each parameter and an initial test run was performed with all parameters at their center value. Next, one test run was carried out for each parameter at their low and high value with all other parameters fixed at their center values. For each test run a variety of descriptive statistics was computed to evaluate the impact of the tested parameter. A precise description of the choices of parameter values and the outcome of the test runs is described in the supplemental materials. Based on these test runs, the default parameter values were all set to their test run center values, except for *q*_min_ whose default value was set to 99.0%.

To demonstrate how the software SDRAP measures up against previous annotation procedures [7, 8], the program was run two more times. Both of the additional tests used the micronuclear genome assembly published in [8] and the macronuclear genome assembly published in [24]. Except for 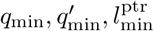 and *k*_max_, all parameters were kept at the default values. To enforce high sequence similarity appropriate for the close relationship between two genomes of the same organism, *q*_min_ and 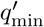 were set to 99% and 95%, for one of the tests and to 95% and 90%, respectively, for the other. Since pointers as short as two base pairs were observed in *O. trifallax*, 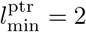 for both tests (see Table 1). To maximize the number of unambiguous subarrangements explored, the parameter *k*_max_ was set to 10 for these test runs. The summary of the runs is shown in Table 2.

**Table 1:** Set of parameter values used for comparison test runs of the software SDRAP. All other parameter values are kept at their default values. The two parameter value sets differ only in the percent identity thresholds for alignments considered in the preliminary and additional match annotation steps.

| Parameter | strict | lenient |
| --- | --- | --- |
| $q_{\min}$ | 99.0 | 95.0 |
| $q'_{\min}$ | 95.0 | 90.0 |
| $l_{\min}^{\text{ptr}}$ | 2 | 2 |
| $k_{\max}$ | 10 | 10 |

**Table 2:**
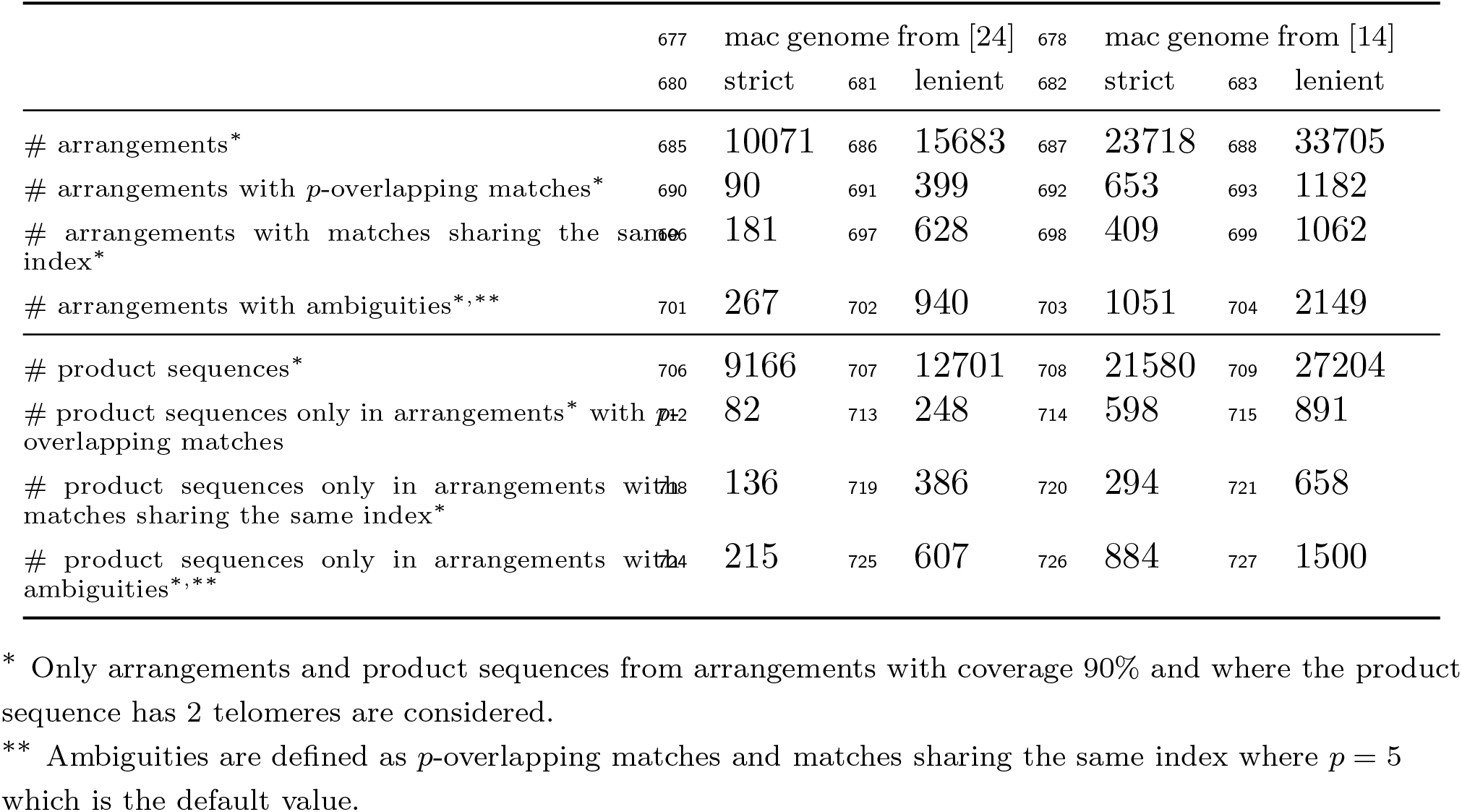
Total counts of arrangements of 2-telomeric product sequences covered at least 90% by the matches in their respective arrangements as well as counts of the subsets of those arrangements that contain ambiguities, along with counts of product sequences only found in these various sets of arrangements. Arrangements were obtained in the two SDRAP test runs applying parameter values listed in Table 1 to the micronuclear genome assembly from [8] and the macronuclear genome assemblies from from [14].

Among the 50 test annotations run on the *O. trifallax* precursor and product genomes *N* is 8,094,251 ± 996,095 (mean ±standard deviation). The number *n*_pre_ of preliminary matches is bounded above by *h*, but among the 50 test runs, the mean number *h* of BLAST alignments across precursor-product sequence pairs with at least one high-scoring pair is 1.7 ± 0.2 (mean ± standard deviation) and the mean ratio *n*_pre_*/h* for these arrangements is 5.5e-2 ±1.3e-1 (mean ± standard deviation). The mean ratio *n*_add_*/h* across arrangements is 3.2e-3 ± 2.6e-3 (mean ± standard deviation). In the 50 test annotations, the mean fraction of arrangements for which the number of unambiguous subarrangements exceeds the threshold out of all arrangements whose scrambling properties were determined is 0.1 ±6.4e-4 (mean ± standard deviation).

Descriptive statistics obtained from the two comparison test runs and corresponding results presented in [8] and [7], respectively, are shown in the supplemental materials. It can be observed that even with the more lenient of the two choices of percent identity thresholds, SDRAP appears to find much fewer matches than [8] (over 15% less matches). However, [8] used ungapped alignments from BLAST whereas SDRAP sometimes merges alignments to form matches. SDRAP obtains a much higher number of arrangements with a single preliminary match. In the more lenient test run, over 45% of these preliminary matches consist of merged matches. Hence, some of the discrepancy between the numbers of single alignment product sequences can be attributed to the artificially introduced fragmentation of alignments in [8] due to the use of ungapped alignments. SDRAP obtains more preliminary matches than were obtained in [7] but only the lenient percent identity thresholds resulted in more matches overall (preliminary and additional). SDRAP detects 36% more scrambled arrangements among strongly complete high coverage arrangements (defined to have coverage at least 90%) than [7]. However, this comparison must be viewed with caution. A different macronuclear genome assembly was used in [7] and the statistics listed for the SDRAP test runs are not necessarily equivalent to their counterparts in [7].

SDRAP was run with the two parameter value sets described in Table 1 using the most recent macronuclear genome assembly published in [14]. The arrangements of 2-telomeric product sequences covered by matches in the arrangement by more than 90% (where coverage is defined as the coverage of the portion of the product sequence between the telomeres) were counted along with the subsets of those arrangements which had *p*-overlapping matches, or matches sharing the same index. These ambiguous rearrangements, or their matches were filtered out in previous procedures. The counts are summarized in Table 2. For the more lenient and stricter choices of parameters, *b*_min_ and *q*_min_ 4.4% and 6.4%, respectively, of all arrangements of 2-telomeric product sequences with coverage at least 90% contained ambiguities in the form of *p*-overlapping matches or matches sharing the same index. Out of all 2-telomeric product sequences that achieved 90% or more coverage in at least one arrangement, 4.1% and 5.5%, respectively, can only be found in arrangements that contain ambiguities. Thus, a more complete annotation of up 6.4% of the data was achieved.

An example of an arrangement that has *p*-overlapping matches is given in Figure 6. The figure visualizes a region of the micronuclear ctg7180000068813 from [8] containing the BLAST high-scoring pairs and precursor intervals matching macronuclear contig AMCR02018712.1 from [14]. Such precise annotation of this arrangement was not obtained with any other previous annotation.

**Figure 6:**
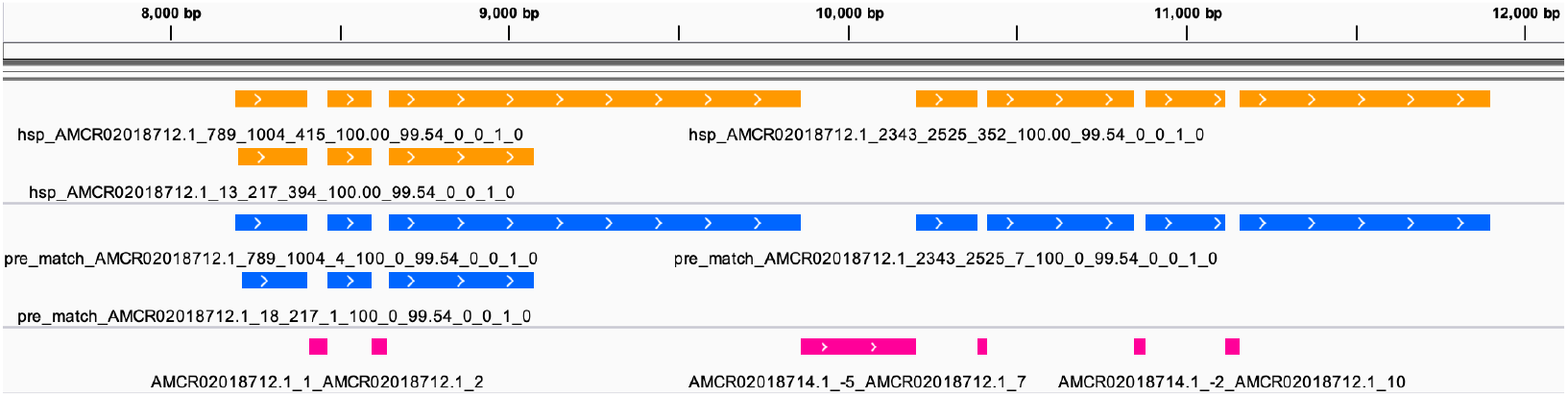
BLAST high-scoring pairs (yellow), matches (blue) and eliminated sequences (pink) on micronuclear sequence ctg7180000068813 from [8]. All alignments and precursor intervals match macronuclear sequence AMCR02018712.1 from [14]. For an explanation of the labelling of the features, see the software documentation at https://github.com/JasperBraun/SDRAP. The picture was obtained using IGV version 2.4.10.

SDRAP was also successfully run on an assembly of PacBio reads from the SK-BR-3 cancer cell line [16] with the PacBio assembly as precursor and the ERBB2 gene region on chromosome 17 in the human reference (GRCh38) as the product. The results confirm extensive duplications and translocations of this locus in the SK-BR-3 cell line, as previously reported by [16], and provide an additional demonstration of the utility of SDRAP for a broad range of data. The output files can be found at https://www.knot.math.usf.edu/SDRAP/annotations/sdrap_skbr3_falcon/.

## 5 Conclusions

Homologous repeats are a prominent issue across most fields that compare DNA sequencing data, regardless of whether the repetitions are inherent to the underlying biology, or an artifact of the procedures, technologies, and protocols used to obtain the data. When analyzing DNA rearrangements, such repetitive regions introduce ambiguity in the establishment of a mapping of precursor to product loci. While various software has been introduced to study DNA rearrangements, to our knowledge, SDRAP is the first that systematically addresses the challenge of many-to-many mappings. Taken together, the present work demonstrates that SDRAP is able both to recapitulate previous workflows and to generate annotations that were not previously possible, while remaining highly flexible and yielding reproducible results. Though SDRAP is especially useful for annotating the genomes of model organisms with programmed DNA rearrangement, it can also enable discovery of genomic rearrangements in other known or unexpected contexts.

## 6 Acknowledgements

Portions of this paper reflect results obtained in the Ph.D. dissertation of author Jasper Braun [5].

## 8 Availability and Requirements

**Project name:** SDRAP

**Project home page:** https://github.com/JasperBraun/SDRAP

**Operating system(s):** Linux

**Programming language:** Javascript, PHP, MySQL

**Other requirements:** PHP 5.3.3, MySQL 5.6.31+, blast+ 2.2.31+

**License:** MIT license

## 9 Abbreviations

SDRAP: Scrambled DNA Rearrangement Annotation Protocol
HSP: High-scoring pair

## 10 Availability of Data and Materials

All datasets used to showcase SDRAP were previously published and are cited in the text. The precursor and product genomes of all runs of SDRAP to annotate rearrangements of *O. trifallax* were obtained from the NCBI Assembly database under assembly names Oxytricha_MIC_v2.0, oxytricha_asm_v1.1, and oxytricha_jrb310_mac_pacbio. Oxytricha_MIC_v2.0 is the precursor assembly used for all runs of SDRAP on the genome of *O. trifallax*. oxytricha_asm_v1.1 is the product assembly for the test and comparison runs. oxytricha jrb310 mac pacbio is the product genome used to generate Figure 6, and both oxytricha asm v1.1 and oxytricha jrb310 mac pacbio were used to generate the data in Table 2.

The annotations generated during the current study are available at (see SDRAP_TESTS_README.tsv). For the precursor of the SK-BR-3 cancer cell line annotation, the Falcon assembly of PacBio reads was downloaded from, and for the product, the ERBB2 locus on chromosome 17 in the human reference GRCh38 was obtained from the NCBI Genome Data Viewer.

## 11 Authors’ Contributions

All authors conceived the project. JB designed and implemented the software. JB and RN conducted and evaluated test runs. All authors participated in manuscript preparation.

