## Supplemental material for "SDRAP for annotating scrambled or rearranged genomes"

This supplement contains parameters and their values used to test SDRAP and present results from these runs.

The parameter $m$, discussed only here, determines the proportion of a product sequence that must be covered by preliminary matches of a precursor sequence for arrangement properties to be computed. This parameter $m$ does not affect how arrangements or arrangement properties are computed. Instead, $m$ only limits how much data is computed, to be able to restrict attention and computational resources to significant arrangements.

In the following tables, $\mathbb{A}_{30}$ and $\mathbb{A}_{90}$ refer to subsets extracted from sets of arrangements ($\mathcal{M}_{\text{pre}} \cup \mathcal{M}_{\text{add}}$, as described in the main manuscript) computed by SDRAP. $\mathbb{A}_{30}$ and $\mathbb{A}_{90}$ represent arrangements where the product sequence without gaps is 30% and 90%, respectively, of the total product sequence (only the product region between telomeric ends, if any, is considered). The notations $\mathbb{A}^*_m$ and $\mathbb{A}^*_{90}$ refer to the sets of arrangements with coverage at least $m$ and 90%, respectively, which did not have more than $k_{\max}$ unambiguous subarrangements. See Table 2 for a reference of these definitions. The output for all test runs can be found at `https://knot.math.usf.edu/SDRAP/annotations/`.

Table 1: Values used for parameters during test runs of SDRAP. For each parameter a "low", "mid", and "high" value was chosen. An initial "center" run with all parameters at their "mid" value was conducted. Only the parameter $l_{\text{min}}^{\text{pre}}$ was set to its "low" value during the initial run. Next, two additional runs for each parameter were performed. Each run differed from the initial test run only in one parameter value which was set to its "low", or "high" value. The parameter $l_{\text{min}}^{\text{pre}}$ was tested at its "mid" and "high" values instead. The "low", "mid", and "high" values for the parameter $\mathcal{S}$ were {ordered}, {ordered, consecutive}, and {ordered, consecutive, complete}. Additional test runs were conducted for some parameters, as indicated in the column "other". The type of each parameter is either SDRAP or BLAST. Parameters labelled type BLAST are used in the implementation in order to filter the alignments provided by BLAST, whereas SDRAP type parameters are specific to the algorithm described in the manuscript and its implementation.

| Parameter | low | mid | high | other | type | description |
|---|---|---|---|---|---|---|
| $l_{\text{min}}^{\text{pre}}$ | 18* | 30* | 35 | | BLAST | Minimum length of alignments for preliminary match annotation |
| $b_{\text{min}}$ | 20 | 49 | 100 | | BLAST | Minimum bitscore of alignments for preliminary match annotation |
| $q_{\text{min}}$ | 60 | 80 | 90 | 95, 99 | BLAST | Minimum percent identity of alignments for preliminary match annotation |
| $c$ | 2 | 4 | 6 | | SDRAP | Minimum contribution to coverage by alignments for preliminary match annotation |
| $s_{\text{max}}$ | 4 | 8 | 12 | | SDRAP | Maximum shift allowed for merging alignments |
| $d_{\text{max}}$ | 1 | 3 | 6 | | SDRAP | Maximum distance between alignments for merging |
| $b'_{\text{min}}$ | 25 | 49 | 100 | | BLAST | Minimum bitscore of alignments for additional match annotation |
| $q'_{\text{min}}$ | 60 | 80 | 90 | | BLAST | Minimum percent identity of alignments for additional match annotation |
| $r$ | 0.5 | 0.8 | 0.9 | | SDRAP | Minimum proportion of product interval alignment must cover to be annotated as additional match and inherit index |
| $r'$ | 0.1 | 0.2 | 0.5 | | SDRAP | Minimum proportion of product interval alignment must cover to be annotated as fragment that inherits index |
| $l_{\text{min}}^{\text{gap}}$ | 2 | 4 | 8 | | SDRAP | Minimum length of non-covered region in product to be considered length |
| $l_{\text{min}}^{\text{ptr}}$ | 1 | 3 | 5 | | SDRAP | Minimum overlap length between successive product intervals to qualify as pointer |
| $\mathcal{S}$ | ** | | | | SDRAP | Collection of properties indicating $\mathcal{S}$-scrambling for unambiguous subarrangements |
| $p$ | 0 | 5 | 10 | | SDRAP | Maximum size of precursor interval overlap tolerated for two matches in an unambiguous subarrangement |
| $k_{\text{max}}$ | 2 | 4 | 10 | | SDRAP | Maximum number of unambiguous subarrangements extracted |
| $m$ | 25 | 50 | 75 | | SDRAP | Minimum proportion of product sequence that must be covered by product intervals of preliminary matches for properties of arrangement to be computed (not detailed in manuscript; used only to limit computation time) |

* $l_{\text{min}}^{\text{pre}} = 18$ was used as the fixed value across test runs for other parameters instead of 30.

** low: {ordered}, mid: {ordered, consecutive}, high: {ordered, consecutive, complete}, additional: {consecutive, complete}

Table 2: Sets of interest in the tables in this supplement.

| set | definition |
|---|---|
| $\mathbb{A}_X$ | subset of arrangements* as described in the manuscript) with coverage** $X$%. |
| $\mathbb{A}^*_X$ | subset of arrangements* with coverage** $x$% and which do not have more than $k_{\max}$ unambiguous subarrangements. |

* An arrangement is defined here as the union $\mathcal{M}_{\text{pre}} \cup \mathcal{M}_{\text{add}}$, as discussed in the main manuscript.

** Coverage refers to the proportion of the product sequence between telomeres, if any, that is not annotated as gapped region.

Table 3: Counts of arrangements and alignments of test runs of parameters related to the MDS annotation step by SDRAP. When a value does not deviate from the value in the row for the center run, it is left blank. The counts of tests of parameters $r$, $r'$, and $l^{\text{ptr}}_{\min}$ do not deviate from the center run and were omitted entirely.

| | | # arrangements ($\mathcal{M}_{\text{pre}} \cup \mathcal{M}_{\text{add}}$) | | | # alignments satisfying $l^{\text{pre}}_{\min}, b_{\min}, q_{\min}$ thresholds** | | # alignments satisfying $b'_{\min}, q'_{\min}$ thresholds** | |
|---|---|---|---|---|---|---|---|---|
| | | total | in $\mathbb{A}_{30}$ | in $\mathbb{A}_{90}$ | in $\mathbb{A}_{30}$ | in $\mathbb{A}_{90}$ | in $\mathbb{A}_{30}$ | in $\mathbb{A}_{90}$ |
| center* | | 429566 | 117612 | 59885 | 1740567 | 891293 | 1740567 | 891293 |
| $l^{\text{pre}}_{\min}$ | mid* | 332994 | 117510 | 59586 | 1720298 | 877743 | 1720298 | 877743 |
| (BLAST) | high | 277861 | 117316 | 59168 | 1686785 | 856651 | 1686785 | 856651 |
| $b_{\min}$ | low | 8167735 | 119159 | 60303 | 1813861 | 923872 | 1744673 | 895206 |
| (BLAST) | high | 184836 | 113585 | 54076 | 1435953 | 711786 | 1710538 | 814542 |
| $q_{\min}$ | low | 446673 | 130942 | 61289 | 1835682 | 927143 | 1772034 | 908144 |
| (BLAST) | high | 366054 | 91517 | 47265 | 1220892 | 647596 | 1535790 | 712381 |
| | 95% | 283489 | 65588 | 27941 | 676173 | 329977 | 1186096 | 400808 |
| | 99% | 160400 | 30010 | 13485 | 269350 | 127346 | 423126 | 168381 |
| $c$ | low | 429582 | 117603 | 59829 | 1740479 | 890265 | 1740479 | 890265 |
| (SDRAP) | high | 429560 | 117612 | 59908 | 1740682 | 891611 | 1740682 | 891611 |
| $s_{\max}$ | low | | 117613 | 59933 | 1740576 | 891874 | 1740576 | 891874 |
| (SDRAP) | high | | 117608 | 59870 | 1740533 | 891153 | 1740533 | 891153 |
| $d_{\max}$ | low | | 117627 | 60010 | 1740707 | 893098 | 1740707 | 893098 |
| (SDRAP) | high | | 117597 | 59855 | 1740387 | 890457 | 1740387 | 890457 |
| $b'_{\min}$ | low | | | | | | 1808171 | 919285 |
| (BLAST) | high | | | | | | 1451418 | 763590 |
| $q'_{\min}$ | low | | | | | | 1782502 | 906120 |
| (BLAST) | high | | | | | | 1294164 | 746688 |
| $l^{\text{gap}}_{\min}$ | low | | 117588 | 59825 | 1740450 | 890390 | 1740450 | 890390 |
| (SDRAP) | high | | 117655 | 60130 | 1740751 | 893959 | 1740751 | 893959 |

* The chosen low value for $l^{\text{pre}}_{\min}$ was used as the fixed value across test runs for other parameters instead of the mid value.

** Numbers of alignments in the columns for $\mathbb{A}_{30}$, $\mathbb{A}_{90}$ are taken from those between precursor and product sequences whose arrangement is in the respective set.

Table 4: Counts of matches for test runs of parameters related to MDS annotation step by SDRAP. When a value does not deviate from the value in the row for the center run, it is left blank. The counts of tests of parameters $r'$, and $l_{\mathrm{min}}^{\mathrm{ptr}}$ do not deviate from the center run and were omitted entirely.

| | | # of preliminary matches | | # of additional matches | | # of merged matches | |
|---|---|---|---|---|---|---|---|
| | | in $\mathbb{A}_{30}$ | in $\mathbb{A}_{90}$ | in $\mathbb{A}_{30}$ | in $\mathbb{A}_{90}$ | in $\mathbb{A}_{30}$ | in $\mathbb{A}_{90}$ |
| center* | | 711568 | 355115 | 407070 | 155486 | 250315 | 149623 |
| $l_{\mathrm{min}}^{\mathrm{pre}}$ | mid* | 703325 | 349741 | 405119 | 154115 | 246390 | 147117 |
| (BLAST) | high | 691449 | 341336 | 400540 | 152568 | 237872 | 144062 |
| $b_{\mathrm{min}}$ | low | 728280 | 361458 | 408032 | 155952 | 254664 | 151778 |
| (BLAST) | high | 605583 | 287584 | 390740 | 146576 | 226193 | 133778 |
| | low | 757659 | 365121 | 404361 | 157424 | 252858 | 151544 |
| $q_{\mathrm{min}}$ | high | 552112 | 292340 | 479820 | 135543 | 256311 | 125787 |
| (BLAST) | 95% | 410835 | 199519 | 446861 | 82703 | 255385 | 66045 |
| | 99% | 242705 | 108125 | 81088 | 28199 | 61009 | 21955 |
| $c$ | low | 714603 | 356373 | 419859 | 159276 | 254675 | 152112 |
| (SDRAP) | high | 709178 | 354278 | 396533 | 153519 | 247391 | 148508 |
| $s_{\mathrm{max}}$ | low | 717737 | 358855 | 402021 | 150824 | 243753 | 144244 |
| (SDRAP) | high | 709400 | 353934 | 407429 | 155907 | 252757 | 151072 |
| $d_{\mathrm{max}}$ | low | 733384 | 366978 | 388375 | 143358 | 217169 | 129691 |
| (SDRAP) | high | 693918 | 346318 | 420928 | 163891 | 275573 | 162754 |
| $b'_{\mathrm{min}}$ | low | | | 412789 | 157385 | 262582 | 152567 |
| (BLAST) | high | | | 359004 | 138948 | 216096 | 133541 |
| $q'_{\mathrm{min}}$ | low | | | 425026 | 159502 | 252549 | 150586 |
| (BLAST) | high | | | 226556 | 107972 | 187068 | 118154 |
| $r$ | low | | | 585559 | 239051 | 306226 | 191833 |
| (SDRAP) | high | | | 341837 | 132860 | 230577 | 136854 |
| $l_{\mathrm{min}}^{\mathrm{gap}}$ | low | 711474 | 354894 | 407068 | 155154 | 250307 | 149424 |
| (SDRAP) | high | 711726 | 356323 | | 155891 | 250331 | 150044 |

* The chosen low value for $l_{\mathrm{min}}^{\mathrm{pre}}$ was used as the fixed value across test runs for other parameters instead of the mid value.

Table 5: Counts of gaps, pointers, and fragments for test runs of parameters related to MDS annotation step by SDRAP. When a value does not deviate from the value in the row for the center run, it is left blank.

| | | # gaps | | # pointers | | # fragments | |
|---|---|---|---|---|---|---|---|
| | | in $\mathbb{A}_{30}$ | in $\mathbb{A}_{90}$ | in $\mathbb{A}_{30}$ | in $\mathbb{A}_{90}$ | in $\mathbb{A}_{30}$ | in $\mathbb{A}_{90}$ |
| center* | | 310040 | 95093 | 401030 | 237895 | 502134 | 264506 |
| $l_{\min}^{\text{pre}}$ | mid* | 312596 | 96650 | 391851 | 231515 | 500301 | 263534 |
| (BLAST) | high | 315783 | 98293 | 379350 | 222514 | 495499 | 259483 |
| $b_{\min}$ | low | 314681 | 94555 | 411635 | 243815 | 503777 | 265273 |
| (BLAST) | high | 305089 | 92489 | 324314 | 182680 | 618642 | 284911 |
| | low | 355106 | 98754 | 420481 | 242795 | 522816 | 272595 |
| $q_{\min}$ | high | 199826 | 67434 | 344141 | 205098 | 404580 | 183182 |
| (BLAST) | 95% | 136408 | 35558 | 270504 | 147755 | 280656 | 69603 |
| | 99% | 60454 | 16813 | 165625 | 79982 | 91449 | 20597 |
| $c$ | low | 309939 | 94911 | 403583 | 239182 | 506619 | 265979 |
| (SDRAP) | high | 310753 | 95562 | 398652 | 236846 | 496788 | 263610 |
| $s_{\max}$ | low | 310043 | 95161 | 402813 | 239024 | 515172 | 276604 |
| (SDRAP) | high | 310044 | 95071 | 400185 | 237384 | 498660 | 261775 |
| $d_{\max}$ | low | 309962 | 95257 | 401836 | 238643 | 556494 | 301596 |
| (SDRAP) | high | 298076 | 90155 | 400119 | 237280 | 463539 | 240079 |
| $b'_{\min}$ | low | | | | | 545623 | 282192 |
| (BLAST) | high | | | | | 443980 | 240485 |
| $q'_{\min}$ | low | | | | | 533474 | 277987 |
| (BLAST) | high | | | | | 382779 | 210390 |
| $r$ | low | | | | | 323645 | 180941 |
| (SDRAP) | high | | | | | 567367 | 287132 |
| $r'$ | low | | | | | 544885 | 296028 |
| (SDRAP) | high | | | | | 393634 | 201707 |
| $l_{\min}^{\text{gap}}$ | low | 322583 | 100747 | 400999 | 237807 | 502123 | 264031 |
| (SDRAP) | high | 279535 | 83193 | 401077 | 238334 | 502140 | 265328 |
| $l_{\min}^{\text{ptr}}$ | low | | | 438964 | 262094 | | |
| (SDRAP) | high | | | 304497 | 178187 | | |

* The chosen low value for $l_{\min}^{\text{pre}}$ was used as the fixed value across test runs for other parameters instead of the mid value.

Table 6: Counts of arrangements for test runs of parameters related to the property computation step by SDRAP. When a value does not deviate from the value in the row for the center run, it is left blank. The total number of arrangements was 429566 for all test runs described in this table. The sizes of $\mathbb{A}_m$ and $\mathbb{A}_{90}$ are 97846, and 59885, respectively, throughout, except for the test runs of $m$. With $m$ at its "low" and "high" value, the two sets have sizes 124066 and 76569, respectively. The counts of tests of parameters $\mathcal{S}$ does not deviate from the center run and was omitted entirely.

| | | # arrangements which exceeded $k_{\max}$ | | # repeating | | # $p$-overlapping | | # repeating and $p$-overlapping | |
|---|---|---|---|---|---|---|---|---|---|
| | | $\mathbb{A}_m \setminus \mathbb{A}_m^*$ | $\mathbb{A}_{90} \setminus \mathbb{A}_{90}^*$ | in $\mathbb{A}_m^*$ | in $\mathbb{A}_{90}^*$ | in $\mathbb{A}_m^*$ | in $\mathbb{A}_{90}^*$ | in $\mathbb{A}_m^*$ | in $\mathbb{A}_{90}^*$ |
| center | | 10380 | 5457 | 7748 | 4835 | 8165 | 4287 | 1550 | 888 |
| $p$ | low | 10453 | 5483 | 7709 | 4824 | 12133 | 6603 | 2538 | 1647 |
| (SDRAP) | high | 10355 | 5450 | 7772 | 4839 | 6924 | 3625 | 1363 | 763 |
| $k_{\max}$ | low | 14102 | 7612 | 4774 | 3040 | 6309 | 3305 | 442 | 266 |
| (SDRAP) | high | 7285 | 3721 | 10039 | 6217 | 10416 | 5508 | 2997 | 1755 |
| $m$ | low | 12198 | | 9262 | | 10002 | | 1845 | |
| (SDRAP) | high | 8031 | | 6110 | | 6327 | | 1248 | |

Table 7: Counts of some arrangement properties for test runs for parameters related to the property computation step by SDRAP. When a value does not deviate from the value in the row for the center run, it is left blank.

| | | # weakly scrambled | | # strongly scrambled | |
|---|---|---|---|---|---|
| | | in $\mathbb{A}_m^*$ | in $\mathbb{A}_{90}^*$ | in $\mathbb{A}_m^*$ | in $\mathbb{A}_{90}^*$ |
| center | | 23534 | 13383 | 17308 | 9730 |
| $\mathcal{S}$ | low | 21198 | 12077 | 15685 | 8968 |
| (SDRAP) | high | 25055 | 14225 | 22554 | 12609 |
| | other | 8580 | 4534 | 7170 | 3675 |
| $p$ | low | 25401 | 14308 | 17970 | 9979 |
| (SDRAP) | high | 22858 | 13047 | 17071 | 9648 |
| $k_{\max}$ | low | 20202 | 11453 | 15874 | 8967 |
| (SDRAP) | high | 26193 | 14850 | 18058 | 10168 |
| $m$ | low | 29531 | | 22102 | |
| (SDRAP) | high | 17867 | | 12960 | |

Table 8: Statistics for the comparison of two SDRAP test runs with results from [2]. A full description of the parameter values used is given in Table 1 of the manuscript. Since the annotation procedures differ between SDRAP and [2], our definitions do not apply directly to the annotations produced by [2], but the attempt was made to define these descriptive statistics as close as possible in meaning to those listed in [2]. Whenever, a directly comparable number could not be obtained due to discrepancies in the annotation procedures, results from [2] are compared to multiple descriptive statistics obtained form the output of SDRAP.

| description | $(q_{\min}, q'_{\min})$ | | reported in [2] |
|---|---|---|---|
| | (99.0, 95.0) | (95.0, 90.0) | |
| # arrangements of 1- or 2- telomeric product sequences with coverage of at least 90% | 12665 | 22844 | 16220 |
| # preliminary matches$^{*}$ | 104069 | 182981 | >225000 |
| # matches$^{*}$ | 105277 | 189513 | |
| # merged preliminary matches$^{*}$ | 861 | 7252 | |
| # alignments merged into preliminary matches$^{*}$ | 1795 | 18244 | |
| # double counted additional matches$^{**}$ | 258 | 1386 | |
| # arrangements that have only one preliminary match$^{*}$ | 1039 | 2896 | 548 |
| # arrangements whose only preliminary match is merged$^{*}$ | 81 | 1309 | |
| # weakly scrambled arrangements$^{*}$ | 1709 | 4140 | 2818 |
| # weakly scrambled arrangements containing matches of both orientations$^{*}$ | 978<br>994 | 2249<br>2355 | 1676 |
| avg # matches over length per product kb in weakly scrambled arrangements$^{*}$ | 4.1/kb<br>4.3/kb | 4.3/kb<br>5.2/kb | 4.9/kb |
| avg # matches per product kb in strongly nonscrambled arrangements$^{*,**}$ | 3.0/kb<br>3.0/kb | 3.0/kb<br>3.0/kb | 3.7/kb |
| median length of preliminary matches in weakly scrambled arrangements$^{*,***}$ | 153bp | 132bp | 81bp |
| median length of preliminary matches in strongly non-scrambled arrangements$^{*,**,***}$ | 196bp | 191bp | 181bp |

$^{*}$ restricted to matches and arrangements of 1- or 2- telomeric product sequences with coverage at least 90%

$^{**}$ excluding arrangements with more than $k_{\max}$ unambiguous subarrangements

$^{***}$ length of preliminary matches does not include pointers

no equivalent to additional matches exists in [2] since repeating MDSs were filtered

Table 9: Statistics for the comparison of two SDRAP test runs with results from [1]. A full description of the parameter values used is given in Table 1 of the manuscript. Since the annotation procedures differ between SDRAP and [1], our definitions do not apply directly to the annotations produced by [1], but the attempt was made to define these descriptive statistics as close as possible in meaning to those listed in [1]. Double counted additional matches are alignments annotated as multiple additional matches due to sufficient overlap with multiple preliminary matches. Such alignments are counted in the numbers of all matches for SDRAP each time they are annotated as additional match.

| description | $(q_{\mathrm{min}}, q'_{\mathrm{min}})$ (99.0, 95.0) | $(q_{\mathrm{min}}, q'_{\mathrm{min}})$ (95.0, 90.0) | reported in [1] |
|---|---|---|---|
| # 2-telomeric contigs | 16027 | | 17198 |
| # product preliminary matches | 422704 | 701725 | 278656 |
| # all matches | 643712 | 1279163 | 752901 |
| # double counted additional matches | 54823 | 190948 | N/A |
| # arrangements with coverage at least 30% | 20713 | 27189 | 39128 |
| # weakly scrambled arrangements with coverage at least 30%* | 2869 | 4176 | 5909 |
| # strongly complete arrangements with coverage at least 90%* | 9925 | 15095 | 15210 |
| # weakly scrambled and strongly complete arrangements with coverage at least 90%* | 1254 | 2109 | 1548 |

* restricted to matches and arrangements of 2-telomeric product sequences which had no more than $k_{\mathrm{max}}$ unambiguous subarrangements
